# Examining the relationship between psychosocial adversity and inhibitory control: an fMRI study of children growing up in extreme poverty

**DOI:** 10.1101/2024.02.05.578942

**Authors:** Zoya Surani, Ted K Turesky, Eileen Sullivan, Talat Shama, Rashidul Haque, Nazrul Islam, Shahria Hafiz Kakon, Xi Yu, William A Petri, Charles Nelson, Nadine Gaab

**Affiliations:** Harvard Graduate School of Education, Cambridge, MA; Laboratories of Cognitive Neuroscience, Division of Developmental Medicine, Department of Medicine, Boston Children’s Hospital, Boston, MA; Infectious Diseases Division, International Centre for Diarrheal Disease Research, Bangladesh; National Institute of Neuroscience and Hospital, Dhaka, Bangladesh; State Key Laboratory of Cognitive Neuroscience and Learning, Beijing Normal University, Beijing, China; Division of Infectious Diseases and International Health, Department of Medicine, School of Medicine, University of Virginia, Charlottesville, VA; Department of Pediatrics, Harvard Medical School, Boston, MA

**Keywords:** Early adversity, functional MRI, executive function, inhibitory control, cognitive development, poverty

## Abstract

Exposure to psychosocial adversity (PA) is associated with poor behavioral, physical, and mental health outcomes in adulthood. As these outcomes are related to alterations in developmental processes, growing evidence suggests that deficits in executive functions–inhibitory control in particular–may, in part, explain this relationship. However, literature examining the development of inhibitory control has been based on children in higher resource environments, and little is known how low resource settings might exacerbate the link between inhibitory control and health outcomes. In this context, we collected fMRI data during a Go/No-Go inhibitory control task and PA variables for 68 children 5 to 7 years of age living in Dhaka, Bangladesh, an area with a high prevalence of PA. The children’s mothers completed behavioral questionnaires to assess the child’s PA and their own PA. Whole-brain activation underlying inhibitory control was examined using the No-Go versus Go contrast, and associations with PA variables were assessed using whole-brain regressions. Childhood neglect was associated with weaker activation in the right posterior cingulate, whereas greater family conflict, economic stress, and maternal PA factors were associated with greater activation in the left medial frontal gyrus, right superior and middle frontal gyri, and left cingulate gyrus. These data suggest that neural networks supporting inhibitory control processes may vary as a function of exposure to different types of PA, particularly between those related to threat and deprivation. Furthermore, increased activation in children with greater PA may serve as a compensatory mechanism, allowing them to maintain similar behavioral task performance.

## INTRODUCTION

Globally, approximately one out of every six adults has been exposed to at least four adverse childhood experiences (ACEs) (Madigan et al., 2023). ACEs are typically described using the categories of abuse, neglect, and household dysfunction, and are used as a measure of the psychosocial adversity (PA) an individual has been exposed to before the age of 18 (Felitti et al., 1998). Individuals who experienced PA in childhood or who were exposed to mothers experiencing PA during this time are found to have increased behavioral, emotional, and academic problems, including increased risk taking, aggressive behavior, and psychopathology, as well as poorer academic achievement (Bick & Nelson, 2016; El-Khodary & Samara, 2020; Lansford et al., 2002). Not surprisingly, exposure to PA produces various physiological and behavioral changes, which can also lead to alterations in developmental processes that can be seen through changes in brain structure, function, and connectivity (Bick & Nelson, 2016; Sheridan et al., 2012; Teicher et al., 2003). There has been recent interest in determining the underlying mechanism that links PA to brain alterations and negative outcomes in adulthood, with one proposed factor involving impaired inhibitory control–a facet of executive functioning (EF) (Wu et al., 2021; Zelazo, 2020; Pechtel & Pizzagalli, 2011). This is an important relationship to consider as understanding the mechanism for which PA relates to psychopathology can better inform interventions in populations with high PA.

### Inhibitory Control

Executive Functions (EFs) comprise a set of discrete cognitive functions including inhibitory control, planning, and cognitive flexibility (Perone et al., 2018). Inhibitory control reflects the ability of an individual to control their impulses by suppressing behavioral habits to make decisions, and can serve as a predictive measure of academic and professional outcomes (Diamond, 2013; Perone et al., 2018; Mann et al., 2017; Kang et al., 2022). In addition to its role in decision-making, inhibitory control is utilized in verbal communication, reading and mathematical skills, memory retrieval, and problem-solving, thus its development in individuals is of particular interest when considering how adulthood outcomes may be affected (Xu et al., 2020).

Inhibitory control has been noted to emerge during infancy, but develops most rapidly between ages 4-6, with further development after this age relating to refinement of inhibitory control networks (Kang et al., 2022; Lee et al., 2015; Liu et al., 2015; Cragg & Nation, 2008). Notable age-related differences from childhood to adulthood include increase in accuracy and speed in task performance and development of brain networks/activation patterns (Cragg & Nation, 2008; Tamm et al., 2002).

When considering brain activation patterns, inhibitory control is represented by a broadly distributed network with the prefrontal cortex as the hub, and more distal sites serving as nodes within this network (Perone et al., 2018). One task that has been commonly used to assess inhibitory control is the Go/No-Go (GNG) task, where participants respond to one stimulus and inhibit their response to a different stimulus. The GNG task is commonly used in fMRI studies, as it is a relatively simple task, where many trials can be completed in a short amount of time to note functional and task-based changes. From fMRI meta-analyses of studies that use the GNG task, brain activation in the prefrontal regions, caudate, cingulate, insula, and parietal cortex were commonly found, primarily in adults (Perone et al., 2018; Simmonds et al., 2008). While inhibitory control has not been extensively studied in young children, previous studies have found greater cortical activation directly related to performance in children compared to adults–reflective of immature prefrontal functioning in children which requires greater activation most likely as a compensatory mechanism (Casey et al., 1997; Lesnik, 2007). Additionally, children’s activation (6-12 years of age) has been observed to be more posterior than adults and less focal (e.g., more widespread area of activation) (Bunge et al., 2002; Lesnik, 2007; Rubia et al., 2003).

It is important to note that the few studies conducted on children have focused on high-income areas, thus the brain correlates of inhibitory control in children growing up in extreme poverty are still unknown. Understanding the neural correlates of inhibitory control in children in low-resourced areas is important, as poverty and factors related to poverty, including PA, can impact inhibitory control development (Pechtel & Pizzagalli, 2011; Wu et al., 2021).

### Relationship between PA Exposure in Childhood and Inhibitory Control

Childhood PA factors have been extensively researched in how they impact children’s growth and development, with inhibitory control commonly assessed alongside to understand how key skills are affected (Felitti et al., 1998; Lund et al., 2020). While exposure to PA has been linked to increased psychopathology, neurodevelopmental models highlight disruption of appropriate EF development, which includes inhibitory control, as a possible mechanism for these outcomes (Zelazo, 2020).

When considering childhood PA factors, particularly maltreatment, most studies have reported greater exposure to PA associated with poorer inhibitory control behavioral outcomes, however some studies reported a lack of significant findings (Carvalho et al., 2020; Lund et al., 2020; Augusti & Melinder, 2013). In light of varying results, rather than considering the cumulative effects of PA as previously done, greater attention is aimed at characterizing and identifying the impacts of the specific forms of adversity as they can manifest in different ways in a developing brain (Felitti et al., 1998; Schäfer et al., 2023). When various types of adversity are assessed as a single factor, developmental differences that may relate to these specific types of PA are unable to be distinguished (McLaughlin, 2016; Schäfer et al., 2023).

For this reason, several investigators have recently proposed that PA can be categorized into two different components: ‘threat’ and ‘deprivation’, where threat includes those that involve exposure to a negative stimulus (e.g., abuse, witnessing or exposure to violence) and deprivation involves those that relate to the absence of an input that is expected in environments to promote development (e.g., neglect) (Schäfer et al., 2023). While threat is expected to have greater associations with impairment in emotional processing, deprivation is more likely related to inhibitory control impairment at a complex cognitive level (Johnson et al., 2021; Schäfer et al., 2023). When considering the nature of the impact of threat, exposure to threat in childhood relates to changes in brain function, particularly the hippocampus and amygdala, due to extensive activation of the hypothalamic-pituitary-adrenal (HPA) axis and maintained detection of heightened stress (Ivy et al., 2010; McLaughlin, 2016; van Marle et al., 2009). Extensive activation of the HPA axis results in increased cortisol secretion, resulting in emotional dysregulation, increased stress, and impaired decision making (Ivy et al., 2010; McLaughlin, 2016; van Marle et al., 2009). These activation changes in the HPA axis can also relate to altered development of the prefrontal cortical circuitry which impacts development of cognitive processes, including those related to EF, e.g., decision making and impulse control (Merz et al., 2023). Deprivation in childhood, however, is more directly associated with cortical changes in the brain, including reductions in dendritic arborization and cortical thickness, mainly impacting the ability to process complex social and cognitive inputs (e.g., those related to EF) (McLaughlin, 2016; Hildyard & Wolfe, 2002). Due to overlap between the categories of adversity, however, the expected outcomes are not exclusive. For example, when considering the relationship between PA and inhibitory control, it is possible that both threat and deprivation result in decreased behavioral performance and reduced brain activation patterns in EF networks, though deprivation would be expected to have greater reductions in these networks activity due to its direct relationship with cortical changes (Johnson et al., 2021).

As a result, from a neuroimaging perspective, this relationship between PA and inhibitory control has yielded mixed results (Kraaijenvanger et al., 2020). Overall, researchers have observed that exposure to various forms of PA has been observed to be associated with decreased activation in key inhibitory control regions, particularly in the anterior cingulate cortex, prefrontal/frontal cortex, and inferior parietal lobule, relating to impaired performance during inhibitory control tasks (Cará et al., 2019; Simmonds et al., 2008; You & Lim, 2015). Additionally, in individuals with similar performance in inhibitory control tasks, differing activation patterns are still observed relating to their PA exposure (Bruce et al., 2013; Jankowski et al., 2017; Ma et al., 2012). More specifically, some researchers have reported decreased activation in task-related areas (e.g., anterior cingulate cortex, middle frontal gyrus) while others have shown increased activation globally, as well as in more posterior regions (e.g., left inferior frontal cortex, right inferior temporal cortex, middle occipital cortex, cerebellum) (Bruce et al., 2013; Ma et al., 2012). It is important to note that as some children exhibit these patterns of increased diffuse activation during the inhibitory control task when there are no significant differences in behavioral performance, one explanation posited is related to compensatory strategies (Ma et al., 2012). It is possible that this greater activation relates to the brain’s ability to increase activity in task-related areas and recruit additional areas to compensate and maintain similar behavioral performance (Ma et al., 2012; Soloveva et al., 2020).

### Relationship between Maternal PA Exposure and Inhibitory Control

Maternal PA, particularly related to mother’s mental health and wellbeing are often used as indicators of stress the child is exposed to in their early environment. In particular, maternal depressive symptoms experienced during early childhood (ages 0 to 6) predicted the child’s difficulties with inhibitory control at later ages (Baker & Kuhn, 2018; Hughes et al., 2013; Wang & Dix, 2017). Reduction of maternal depressive symptoms were also observed to be associated with restoration of EF deficits, i.e. higher EF (Lund et al., 2020). Additionally, positive parenting practices (e.g., warmth and sensitivity toward children) have been associated with lower inhibitory control network connectivity–which is associated with maturation of higher-order networks (Pozzi et al., 2021). It is hypothesized that exposure to maternal stress not only impacts the HPA axis resulting in increased cortisol secretion (i.e., stress response) and impacts to prefrontal cortical circuitry, but as it can be related to lower levels of maternal sensitivity and care, it can impact social and cognitive networks more directly as well (e.g., those related to inhibitory control) (Gonzalez et al., 2012; Bernier et al., 2010; Saridjan et al., 2010; Merz et al., 2023).

Although the impact of depression on children’s inhibitory control has been examined at a behavioral level, the impacts on children’s neural substrates of inhibitory control remain largely unexplored, particularly in low-income areas. Research conducted outside of the early childhood years is unable to consider the early brain networks that occur as the preschool period is a period of rapid inhibitory control maturation (Lee et al., 2015; Liu et al., 2015). Finally, understanding how this relationship manifests in low-resource environments is largely necessary to understand how this relationship manifests in areas with a high prevalence of PA. In a recent study that compared the neural correlates of inhibitory control through EEG and ERP in Dhaka, Bangladesh as part of the Bangladesh Early Adversity Neuroimaging (BEAN) study, it was found that children with greater early adversity had lower accuracy, lower IQ, and notable neural differences (e.g., more pronounced P3 amplitude in the inhibitory control condition) (Sullivan et al., 2022). As differences were found in this population using EEG and ERP neuroimaging tools, inherently, it is necessary to understand these changes through additional functional methods (e.g., fMRI).

### The Current Study

The purpose of the current study is to extend previous research to examine the relation between PA and the functional neuroanatomy of inhibitory control, specifically using fMRI data collected during the GNG task, in children 5 to 7 years of age in an environment with a high prevalence of extreme poverty in Dhaka, Bangladesh. In this area, 64% of families met the World Bank definition of extreme poverty, which includes families living on less than $1.90 per day (Sullivan et al., 2022). These are important considerations, not only to understand if existing trends from research done in high- and middle-resource countries extend to low-resource countries where there is a greater prevalence of PA, but also to examine the brain function underlying the differences in inhibitory control among children at an age where inhibitory control is developing rapidly. Understanding how this relation manifests in an area of high poverty, particularly from a neuroimaging level, will allow for greater identification of the alterations that occur in developmental processes associated with exposure to PA.

We sought to examine this relation and add to existing research through three main aims. First, we examined the neural basis of inhibitory control through fMRI in children living in a low-resource environment to determine whether previous activation patterns found are generalizable to our population. Despite the recent growth and progress in the developmental cognitive neuroscience field, few studies have been conducted in cultural and contextual environments that extend from the traditional Western, educated, industrialized, rich, and democratic (WEIRD) societies (Garcini et al., 2022). By examining how activation patterns from our sample compare to those observed in prior research (e.g., activation in the prefrontal cortex, cingulate; widespread areas of activation in superior and middle frontal gyri), a new perspective of brain development will be added.

Second, we assessed whether inhibitory control activation varied with exposure to PA. Specifically, we performed regression analyses to determine how activation patterns in the inhibitory control contrast were related to children’s exposure to specific types of PA as well as a cumulative measure (combining the individual PA factors). Finally, we determined how activation patterns for inhibitory control differ between children with varying exposure to maternal PA. When considering the role of PA, we hypothesize that children who experienced greater PA, childhood and/or maternal, particularly in the cumulative measures, would have inefficient recruitment of task-related areas, including decreased activation in key inhibitory control areas, including the parietal and prefrontal cortices, as well as the anterior cingulate, as shown in Kraaijenvanger et al. (2020), Cará et al. (2019), and Bruce et al. (2013). When considering specific PA factors, we hypothesize that those related to deprivation (e.g., neglect, social isolation) would more likely have greater observable differences in activation patterns than those related to threat due to the nature of the cognitive task (Johnson et al., 2021; Schäfer et al., 2023). As a compensation mechanism for individuals with similar performance but differing levels of PA, we hypothesize that there may be increased activation globally as observed by Bruce et al. (2013) and Ma et al. (2012). Successful completion of these goals will help characterize the neural correlates of inhibitory control in children in a low-resource environment and identify how activation between EF-related regions is altered in children with exposure to PA.

## METHODS

### Participants

The participants’ data used for this project are from a larger study, the Bangladesh Early Adversity Neuroimaging (BEAN) Study. The BEAN study aims to understand the relation between biological adversity and psychosocial adversity (PA) and neurocognitive development in children in Dhaka, Bangladesh. Participants were recruited from a medical clinic in the Mirpur area of Dhaka. All children included in the study were born at greater than 34 weeks gestation and had no history of neurological abnormalities/traumatic brain injury, known genetic disorders, and/or visual or auditory delays or impairments (Jensen et al., 2019).

Children were drawn from two cohorts: the Performance of Rotavirus and Oral Polio Vaccines in Developing Countries (PROVIDE) or the Cryptosporidium Burden Study (CRYPTO) cohorts. Children from the PROVIDE and CRYPTO cohorts were initially recruited as two separate groups, but as all children were from a low-resource environment, whose PA and fMRI data were collected in similar ways, their PA and MRI data were processed identically and combined into one group.

Whereas 260 children were included in the larger BEAN study, the current study includes a smaller subset due to challenges in fMRI and psychosocial questionnaire data collection as well as exclusion criteria used for the fMRI data (related to motion and in-scanner performance), further explained in fMRI Data Processing. The final sample of combined CRYPTO and PROVIDE cohorts included 68 children (34 M, 34 F), with an average age of 78.9 months (SD = 6.4, range = 61 – 84).

This study received ethical approval from research and ethics review committees at The International Centre for Diarrhoeal Disease Research, Bangladesh (ICDDRB), and the Institutional Review Board at Boston Children’s Hospital.

### Psychosocial Adversity Measures

When participants were 60 months of age, trained research assistants interviewed participants’ mothers in order to complete a comprehensive behavioral battery to assess various PA measures, including the Childhood Psychosocial Adversity Scale (CPAS) (Berens et al., 2019), Edinburgh Postnatal Depression Scale (EPDS) (Gausia et al., 2007), Perceived Stress Scale (PSS) (Cohen et al., 1983), and Tension Scale (TS) (Karasz et al., 2013). The CPAS was used to assess the children’s cumulative exposure to PA while the EPDS, PSS, and TS were used to assess the presence of maternal PA. Descriptive statistics with example questions for each questionnaire are presented in Table 1.

**Table 1:**
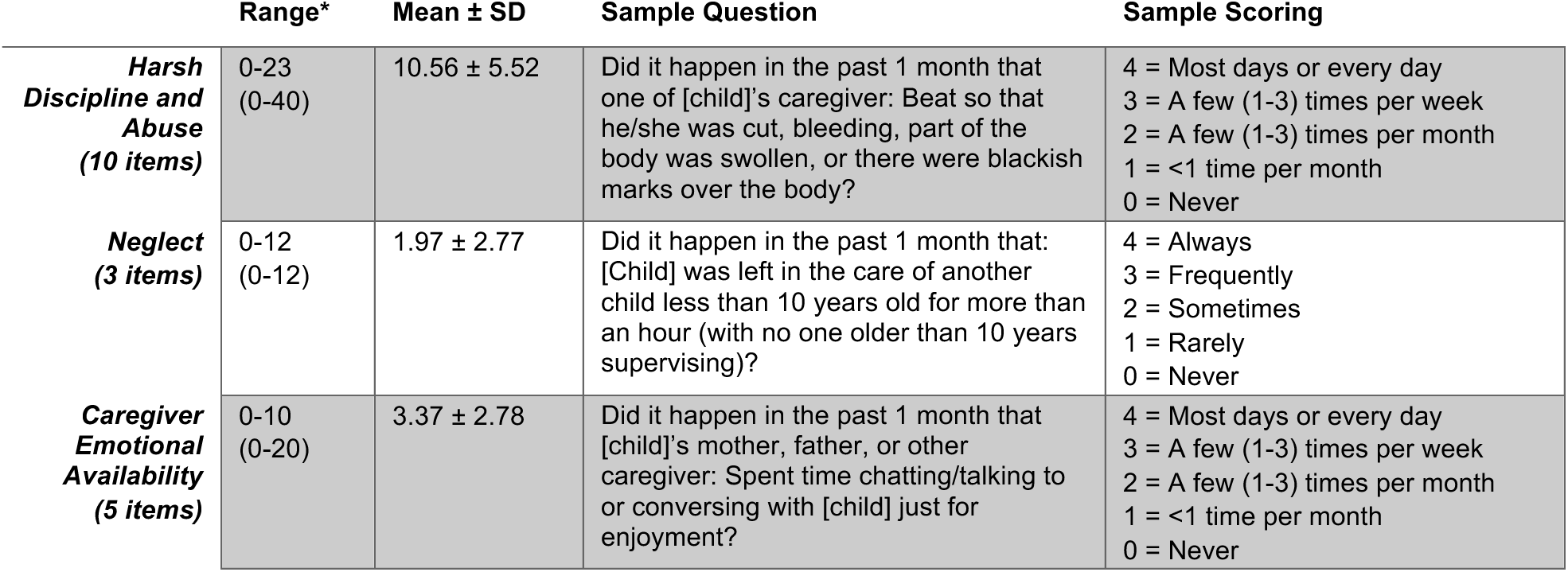
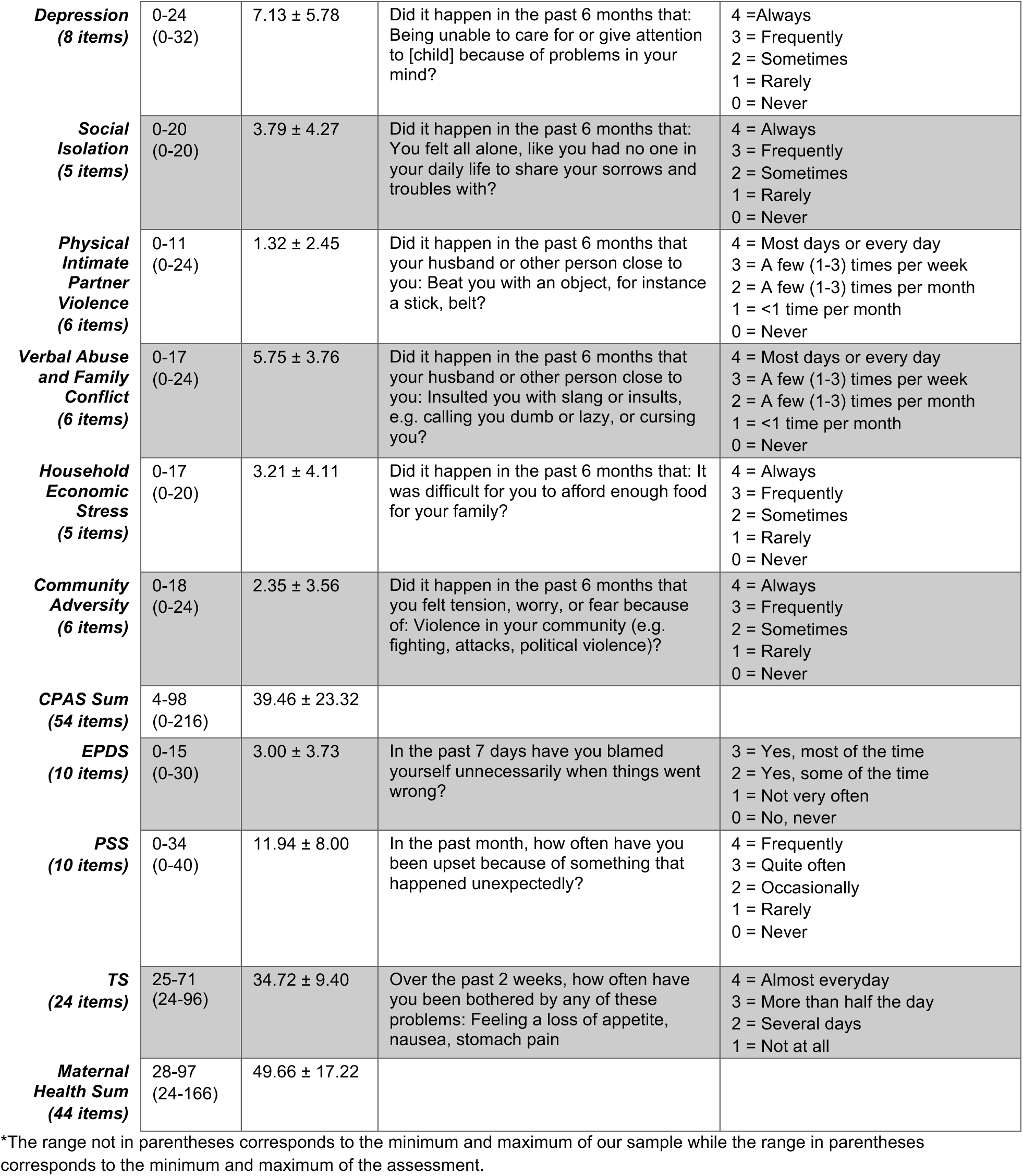
Descriptive measures for childhood and maternal PA measures.

#### Childhood PA

The CPAS was a locally adapted parent-reported questionnaire designed specifically for this study sample through qualitative work with Bangladeshi mothers, field staff, and psychologists, and pretested in the current study sample prior to use. The CPAS included 54 total items rated on a scale of 0 (never) to 4 (always/most days or every day) grouped into nine subscales with varied numbers of items in each, to determine the child’s exposure to different forms of adversity by assessing the frequency of adverse events (Berens et al., 2019). The CPAS subscales (Harsh Discipline and Abuse, Neglect, Caregiver Emotional Availability, Depression, Social Isolation, Physical Intimate Partner Violence, Verbal Abuse and Family Conflict (Family Conflict), Household Economic Stress, and Community Adversity) were used as individual measures and as a simple sum measure (CPAS Sum) according to the scale’s indicated grading (Berens et al., 2019). The Caregiver Emotional Availability subscale was reverse scored for this sum value as it assessed positive characteristics of the home environment. The subscales were found to be highly interrelated (Cronbach’s α = 0.82). The average childhood PA summary score was 39.46 ± 23.32 (of a max score of 216).

#### Maternal PA

In addition to the CPAS, the EPDS, PSS, and TS, were collected to assess maternal mental health and wellbeing. The EPDS was previously validated in Bangladesh and had 10 items rated on a scale from 0 (never) to 3 (most of the time) (for a max score of 30) that related to the mother’s mood and feelings for the past 7 days in her postpartum period (Gausia et al., 2007). Two of the items were reverse coded to ensure higher scores reflect higher levels of maternal stress. The PSS, which is commonly used to capture a mother’s thoughts/feelings in the past month, consisted of 10 items rated on a scale from 0 (never) to 4 (very often), with 3 items reverse coded. The PSS was translated, back-translated, and pretested in Bengali with experts and field populations as part of the BEAN study (Sullivan et al., 2022). The TS, which assesses the frequency of various emotional and physical manifestations of stress over the past 2 weeks, had 24 items rated on a scale of 1 (not at all) to 4 (almost every day) (for a max score of 96). The TS was designed specifically for this population (i.e., Bangladeshi women) and was used without any changes (Karasz et al., 2013). The three questionnaires were used as individual measures as well as a simple sum score to analyze the relationship between specific types of PA in addition to a total PA exposure value. The three questionnaires were found to be highly interrelated in the larger BEAN sample (Cronbach’s α = 0.72), while our sample was found to have an interrelation of 0.66 (Sullivan et al., 2022). The average maternal PA summary score was 49.66 ± 17.22 (of a max score of 166).

### Go/No-Go Task Paradigm

An event-related Go/No-Go (GNG) task was used to identify brain correlates of inhibitory control, one facet of executive functioning. This task was modeled after the GNG task used by Langer et al. (2019), but as Bangladeshi children were less familiar with the images of crabs initially used as stimuli, these images were replaced. Children were instead presented with either an image of a rooster or an image of a chick and told to respond as quickly as possible when the rooster appeared (Go condition) and refrain from responding when the chick appeared (No-Go condition). The task included a practice run, which occurred outside the scanner to ensure the participants understood the task design, and three experimental runs in the scanner. The practice run had a minimum of 30 trials and variable percentages of “Go” and “No-Go” stimuli between participants, while the three experimental runs each contained 80 trials with 75% of the stimuli presented being “Go” stimuli and 25% of the stimuli presented being “No-Go” stimuli. The “Go” stimuli did not appear more than three times in a row, and the “No-Go” stimuli did not appear more than two times in a row. Additionally, each run began with a “Go” stimulus. Each stimulus was displayed for 1000 ms and was followed by an interstimulus fixation interval. The interstimulus fixation screen was displayed for between 1000 ms and 9000 ms and participants could respond at any point during this interval by pushing (or refraining from pushing) a button on a simple button remote. Variable interstimulus intervals were used to decrease stimulus anticipation (Condy et al., 2020; Wodka et al., 2009). While the trial-to-trial variability of the interstimulus intervals were randomized during task design, once complete, all participants received the same task, with the same ordering of interstimulus intervals. A representation of the task design is included in Figure 1.

**Figure 1:**
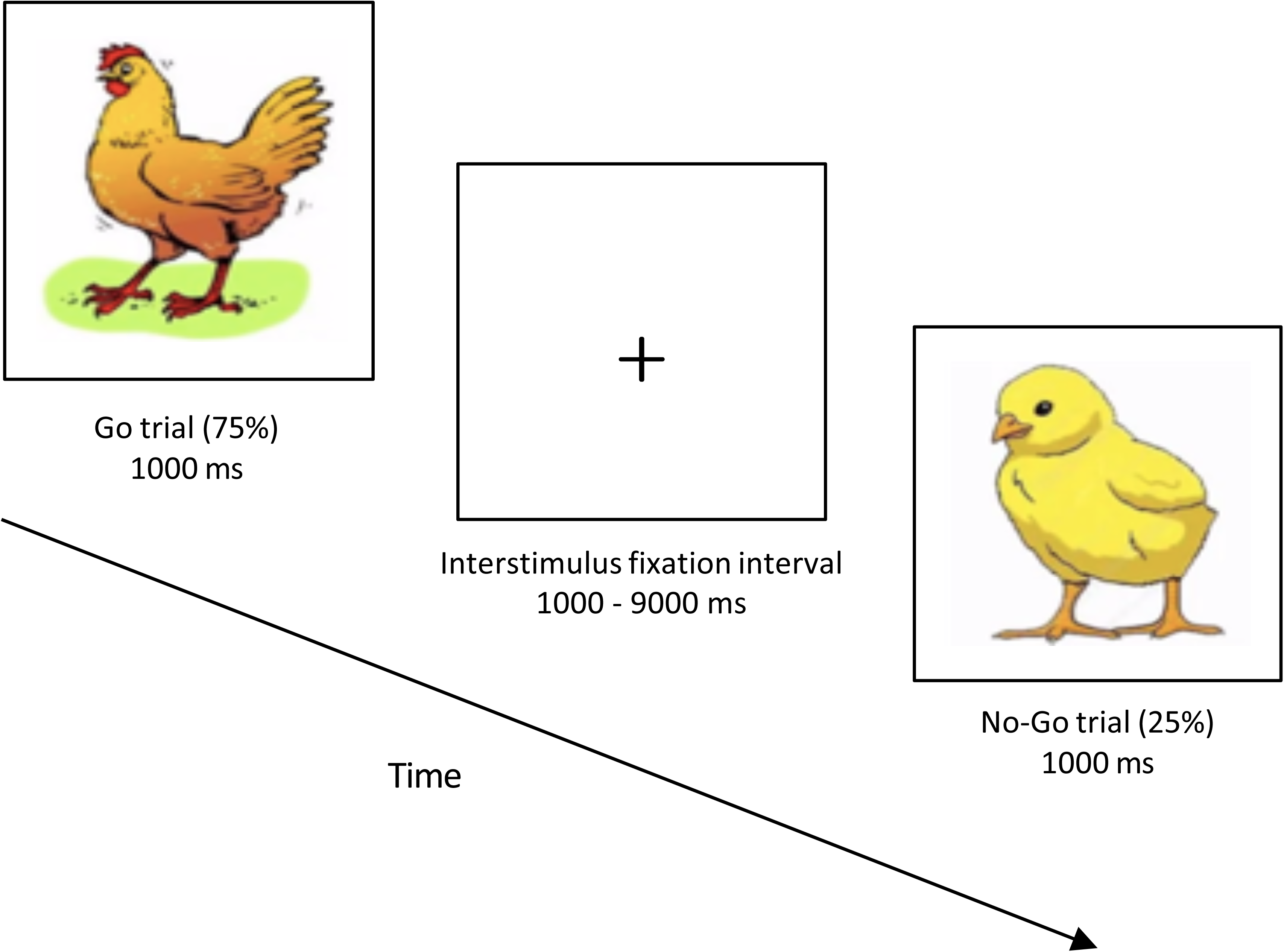
Go/No-Go task design. This figure depicts the task design for the GNG task used. The stimulus presentation was followed by the presentation of a fixation screen (which varied in length) in which the participant responded. After the fixation interval, a new stimulus would appear. When the rooster stimuli appeared, children were asked to respond by pressing a button, whereas when the chick stimuli appeared, children were asked to refrain from responding.

### MRI Data Acquisition

Prior to the start of MRI scanning, local staff from the National Institute for Neuroscience and Hospital (NINSH) and ICDDRB in Dhaka, Bangladesh visited Boston Children’s Hospital to receive training for conducting pediatric MRI. While the center had previously scanned pediatric patients for clinical and pilot studies, this training ensured staff were familiar with the requirements needed for this large-scale study. A protocol was designed that details this training and the limitations for conducting MRI in low-resource settings (Turesky et al., 2019).

As explained in Turesky et al. (2021), participants were scanned in a 3T Siemens MAGNETOM Verio scanner using a 12-channel head coil at the NINSH. Consent was obtained the day prior to the children’s scanning. Before the children were scanned, they were shown various anatomical images of the brain to understand the function of the MRI scanner and were also able to familiarize themselves with a mock cardboard scanner and set of button controls. Information related to the parameters of the scans are included in the Supplementary Material (Table 1).

### fMRI Data Processing

#### Preprocessing

Image preprocessing, including slice-time correction, realignment, coregistration, normalization, smoothing, and artifact removal, was completed in SPM12. Images were realigned to correct for head movement over the course of the scan and any images with large spikes in head movement were removed. Normalization was performed to ensure brain images were in the same space. Smoothing (full-width at half maximum = 8mm) was done to improve the signal-to-noise ratio. Participants for whom 30% of their volumes were preceded by over 2mm of head motion were removed from further analysis and the functional dataset.

#### Analysis

For all participants who met accuracy (above 50% average) and motion requirements, the three experimental runs of the GNG tasks were combined in a fixed-effects model. A high-pass filter (128 Hz) was used to remove low-frequency noise before creating a multiple regression model. Regression analyses as part of participant-level statistics were conducted on the processed functional time series to examine the activation over the time course. Separate models were created for the correct trials of the Go condition (Go stimuli pressed) and No-Go condition (No-Go stimuli unpressed). Contrasts were generated for Go > Fixation (baseline), No-Go > Fixation, and No-Go > Go (NGvG) in individual subjects to compare activity during the task against their baseline. Fixation represented the participant’s baseline, or resting, activation, which was measured during the fixation portion of the GNG task. The NGvG contrast was generated to visualize which regions underlie inhibitory control.

Group statistics involved combining individual brain maps to create group maps to visualize regions active in these contrasts. The height threshold used for the contrasts was p = 0.001. The FDR cluster-level corrected p-value was 0.05. Clusters of activation, peak intensities, peak coordinates, and loci in MNI space were ascertained using SPM12. MNI coordinates were then transformed into Talairach anatomical space (Talairach et al., 1988) using the icbm2tal algorithm within the GingerALE platform (Lancaster et al., 2007), and anatomical regions were labeled according to the Talairach Daemon (http://www.talairach.org/daemon.html).

### Brain-Behavioral Analyses

To understand the relation between PA and children’s inhibitory control, a data-driven approach was used, involving multiple regressions run on a whole-brain level. Regressions were run on the NGvG contrast with the individual childhood PA measures (CPAS subscales), the individual maternal PA measures (EPDS, PSS, TS), and the sum childhood PA (CPAS Scaled) and maternal PA (Maternal Health Sum) scores listed as covariates of interest in separate models. Given that two cohorts were combined for these analyses, a covariate of no interest was added to distinguish the PROVIDE and CRYPTO cohorts.

Brain-behavioral analyses were used to determine if there was a statistically significant relationship between the PA variable and NGvG activation. Both positive and negative contrasts were examined to ensure that the direct and inverse relationships between each PA measure (14 scales) and activation subserving inhibitory control were considered, resulting in a total of 28 models. Within each model, false discovery rate correction was applied to correct for the number of voxels in the brain, whereby each voxel represents a separate statistical test. Standard practice in fMRI studies does not typically include correction for the number of whole-brain models tested, as corrections within each model are already highly stringent due to the number of voxels (tens of thousands) (Bruce et al., 2013; Turesky et al., 2019). The height threshold used for brain-behavioral analyses was p = 0.001. The FDR cluster-level corrected p-value was 0.05.

## RESULTS

### Behavioral Results

After combining the accuracy values for participants (n = 68) in the three Go/No-Go experimental runs, the average accuracy for the Go and No-Go runs were calculated. Similar to results from previous studies with this age group, the accuracy for the Go runs (89.7%, *SD = 6.9*) was higher than for the No-Go runs (83.5%, *SD = 12.0*) (p < 0.001) (Sullivan et al., 2022; Torpey et al., 2012). The accuracy values exclude participants who performed with less than 50% accuracy (n = 7), as these participants were removed from the final analysis.

Correlations were run between the accuracy values (Go accuracy, No-Go accuracy) with childhood and maternal psychosocial adversity (PA) variables, as well as participant age. No significant correlations were found between any of these measures. Significant correlations were found between various individual childhood and maternal PA variables. These results are included in Supplementary Material (Table 2).

**Table 2:**
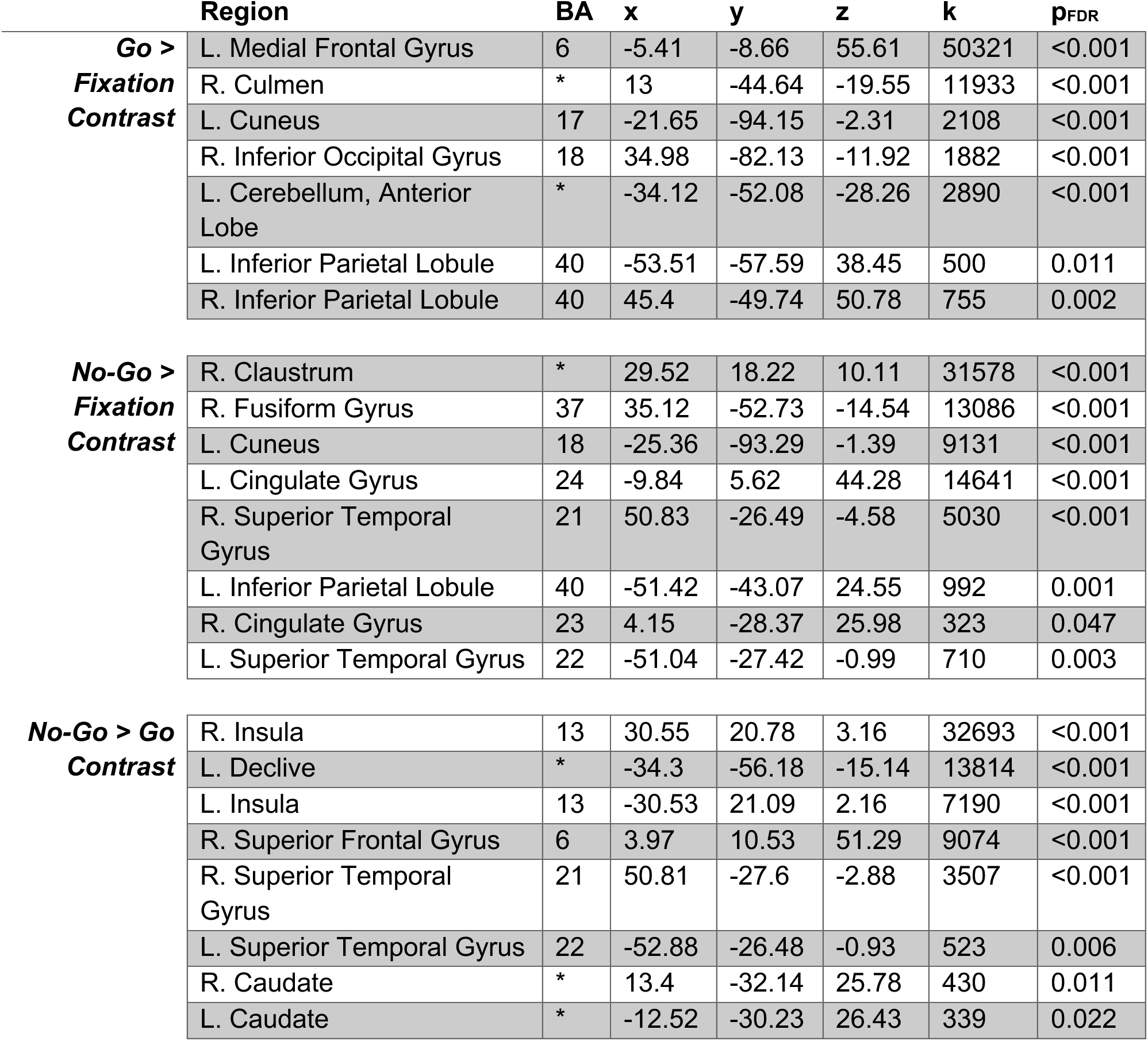
Locations of significant whole-brain activation cluster peaks for the Go, No-Go, and No-Go > Go contrasts.

### Neuroimaging Results

#### Go/No-Go Contrasts: Group Maps

Whole-brain maps were created for the Go > Fixation, No-Go > Fixation, and No-Go > Go (NGvG) contrasts. Table 2 includes the coordinates and details for each of the significant cluster peaks from the three contrasts. Figure 2 depicts the surface renderings of these areas.

**Figure 2:**
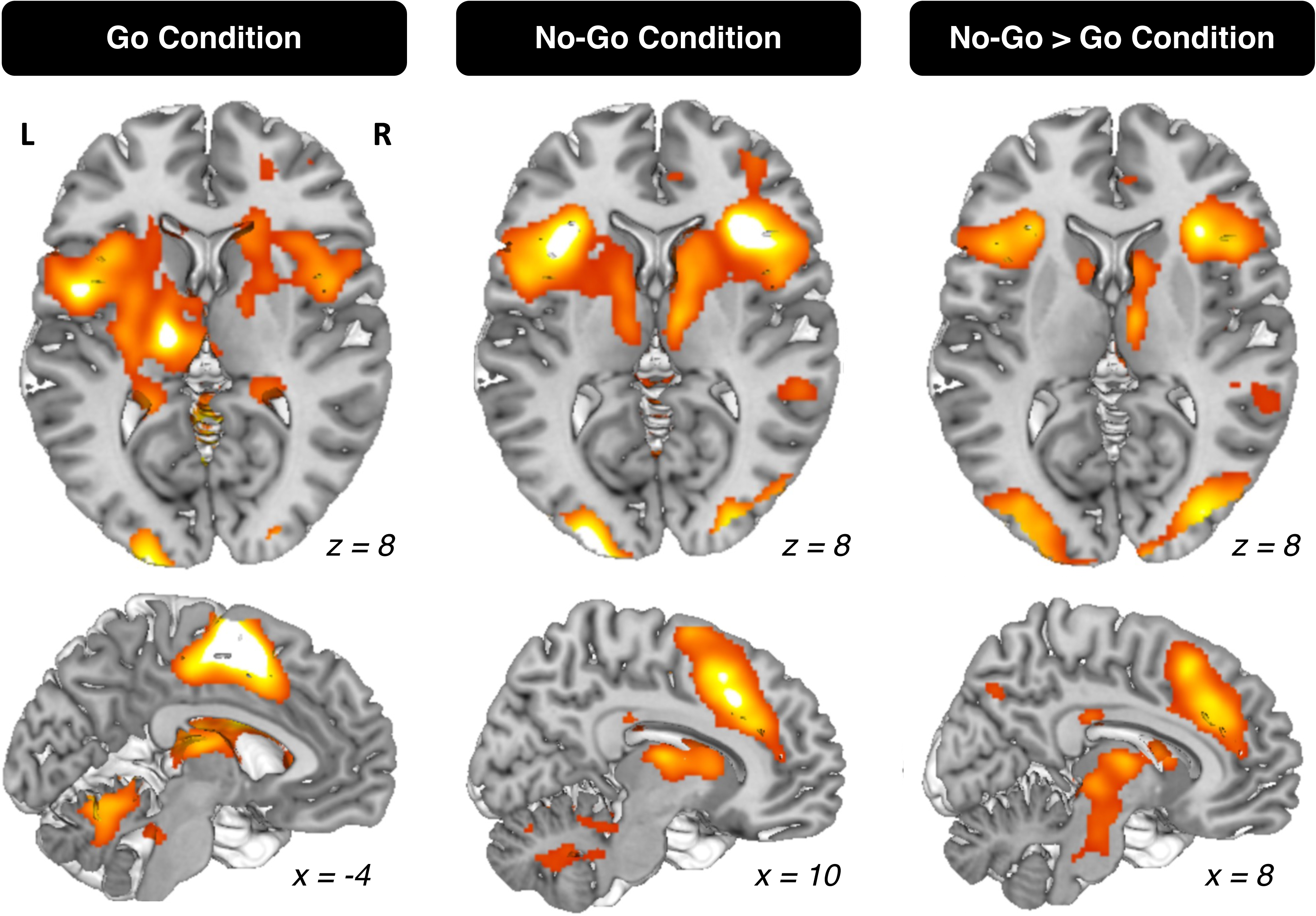
Activation maps for Go, No-Go, and No-Go > Go Contrasts. Whole-brain activation maps for the Go, No-Go, and No-Go > Go contrasts in the GNG task (FDR cluster-level correction of p < 0.05). Activation was found through whole-brain regression group analyses (n = 68). Surface renderings of activation patterns were created in Mango (http://ric.uthscsa.edu/mango/) with significant clusters overlayed on Colin27 T1 segmented MNI template. Views shown: superior axial (top) and sagittal right (bottom). L and R represent the left and right sides of the axial surface, respectively. The x and z values represent the position of the respective axes from which the image was taken (x = left/right, z = top/bottom). Table 2 provides the full list of activations revealed by each of these contrasts.

For the Go > Fixation contrast, significant activation clusters were found in the left medial frontal, and bilateral occipital, cerebellar, and inferior parietal regions. The peak in the large left medial frontal cluster extended into the right medial frontal, left pre- and postcentral gyri, and bilateral basal ganglia, thalamus, and insula areas. The No-Go > Fixation contrast revealed significant activation clusters in the right claustrum, left inferior parietal, and bilateral occipital, cingulate, and superior temporal areas. The peak in the right claustrum cluster extended into the bilateral insula, basal ganglia, and thalamus. The NGvG contrast revealed significant activation in the right superior frontal, left cerebellar, and bilateral insula, superior temporal, and basal ganglia areas. Right cerebellar, and bilateral cingulate and occipital regions were also active through extensions from other areas.

#### Psychosocial Adversity: Childhood PA

To examine whether greater childhood PA was associated with weaker activation (i.e., inverse relationship), whole-brain regressions were run between the NGvG contrast across the whole brain and CPAS subscales and summary scores. The NGvG contrast was used for these brain-behavior analyses because the activation patterns in this condition represented those that subserve inhibitory control (Simmonds et al., 2008). It was hypothesized that these associations would be greater in subscales related more to deprivation (e.g., neglect, caregiver emotional availability, social isolation).

Greater childhood neglect was associated with weaker NGvG activity in the right posterior cingulate (p_FDR_ = 0.014; peak coordinates: x = 1.42, y = −61.87, z =12.85; k = 515 voxels). Neither the childhood PA sum score nor any other childhood PA subscale were negatively associated with NGvG activity.

To examine whether greater PA was associated with greater activation, possibly due to compensatory activation patterns as there were no differences in task performance observed, whole-brain regressions were used to test the direct relationship between childhood PA and the NGvG activation through a positive contrast.

Greater childhood family conflict was associated with greater activation in the left medial frontal gyrus extending to the left anterior cingulate (p_FDR_ <0.001; peak coordinates: x = −8.8, y = 28.41, z = 41.95; k = 2068 voxels), while greater household economic stress was associated with greater activation in the right superior frontal gyrus extending to the right medial frontal gyrus (p_FDR_ = 0.019; peak coordinates: x = 9.35, y = 1.92, z = 62.27; k = 601 voxels). Brain maps depicting surface renderings of the activation present in both the negative and positive contrasts are provided in Figure 3.

**Figure 3:**
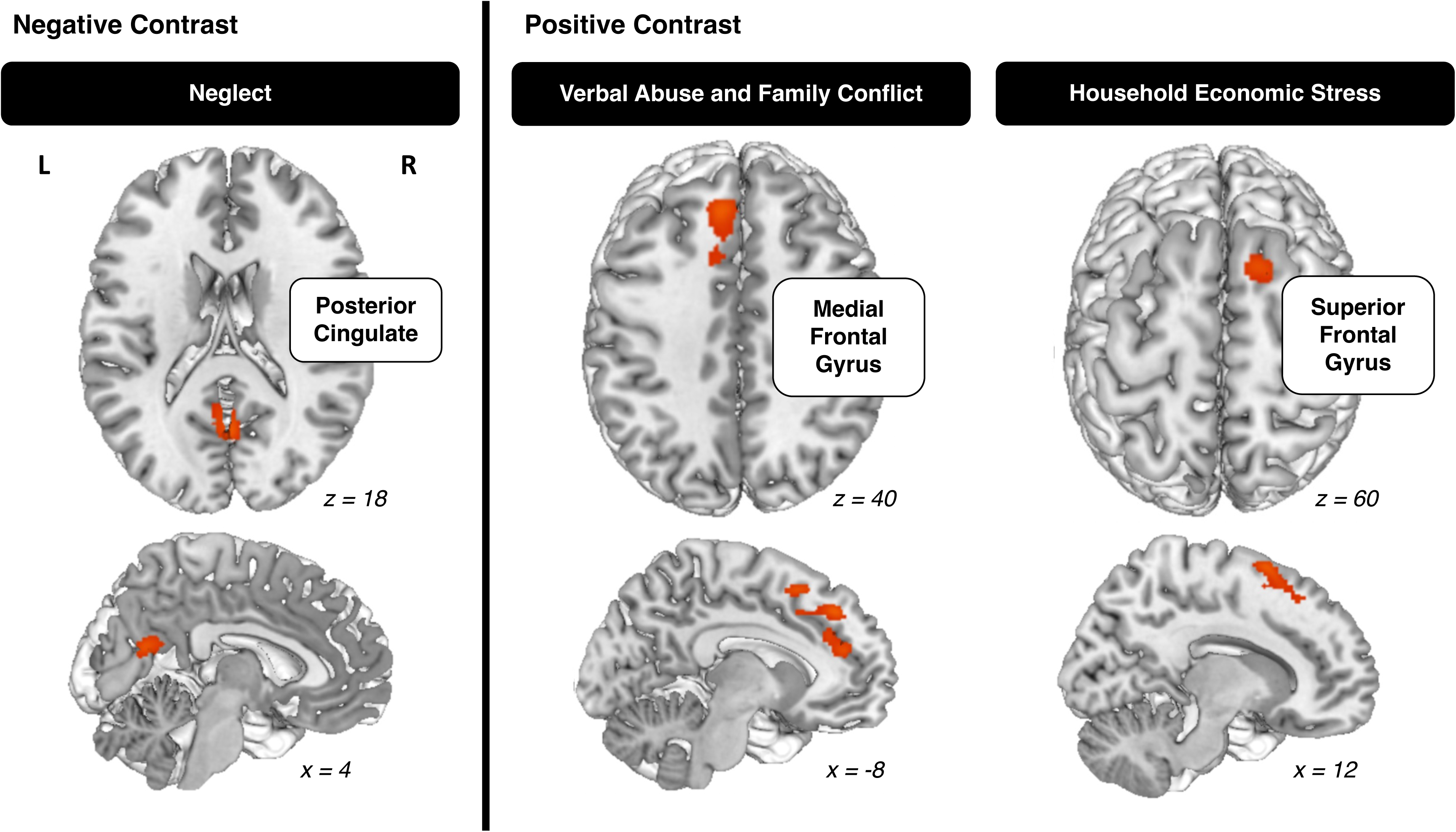
Locations of significant whole-brain activation clusters for No-Go > Go contrast (FDR cluster-level correction of p < 0.05) with childhood psychosocial adversity measures as covariates for both positive and negative contrasts. Activation was found through whole-brain regression group analyses (n = 68). Surface renderings of activation patterns were created in Mango (http://ric.uthscsa.edu/mango/) with significant clusters overlayed on Colin27 T1 segmented MNI template. Views shown: superior axial (top) and sagittal right (bottom). L and R represent the left and right sides of the axial surface, respectively. The x and z values represent the position of the respective axes from which the image was taken (x = left/right, z = top/bottom).

#### Psychosocial Adversity: Maternal PA

Similar to the reasoning used for the childhood PA analyses, whole-brain regressions were run with both negative and positive contrasts between the NGvG contrast and the maternal PA questionnaires (i.e., mothers reporting their own experiences of PA) and summary score included as covariates.

For all regression models–the three individual maternal health questionnaires (EPDS, PSS, TS) and summary score, an inverse relationship of activation from the NGvG contrast, where higher maternal PA values correlate with weaker NGvG activity, was not found.

In the positive contrast, significant clusters were found for all covariates, with greater maternal PA assessed through the EPDS and PSS associated with increased activation (EPDS: p_FDR_ = 0.001, k = 1100 voxels; PSS: p_FDR_ = 0.005, k = 765 voxels) in the left cingulate gyrus extending into the left medial and superior frontal gyri (peak coordinates: x = −10.69, y = 24.43, z = 44.25). Greater TS was associated with increased activation in the right middle frontal gyrus extending into the right cingulate gyrus (p_FDR_ = 0.015; peak coordinates: x = 37.39, y = 11.14, z = 43.8; k = 665 voxels), and the overall summary score associated with increased activation in the left medial frontal gyrus extending into the left anterior cingulate (p_FDR_ <0.001; peak coordinates: x = −8.8, y = 28.41, z = 41.95; k = 2298 voxels).

Figure 4 depicts the results and surface renderings of the positive contrast. As there were no significant results from the negative contrast, there are no maps for that contrast.

**Figure 4:**
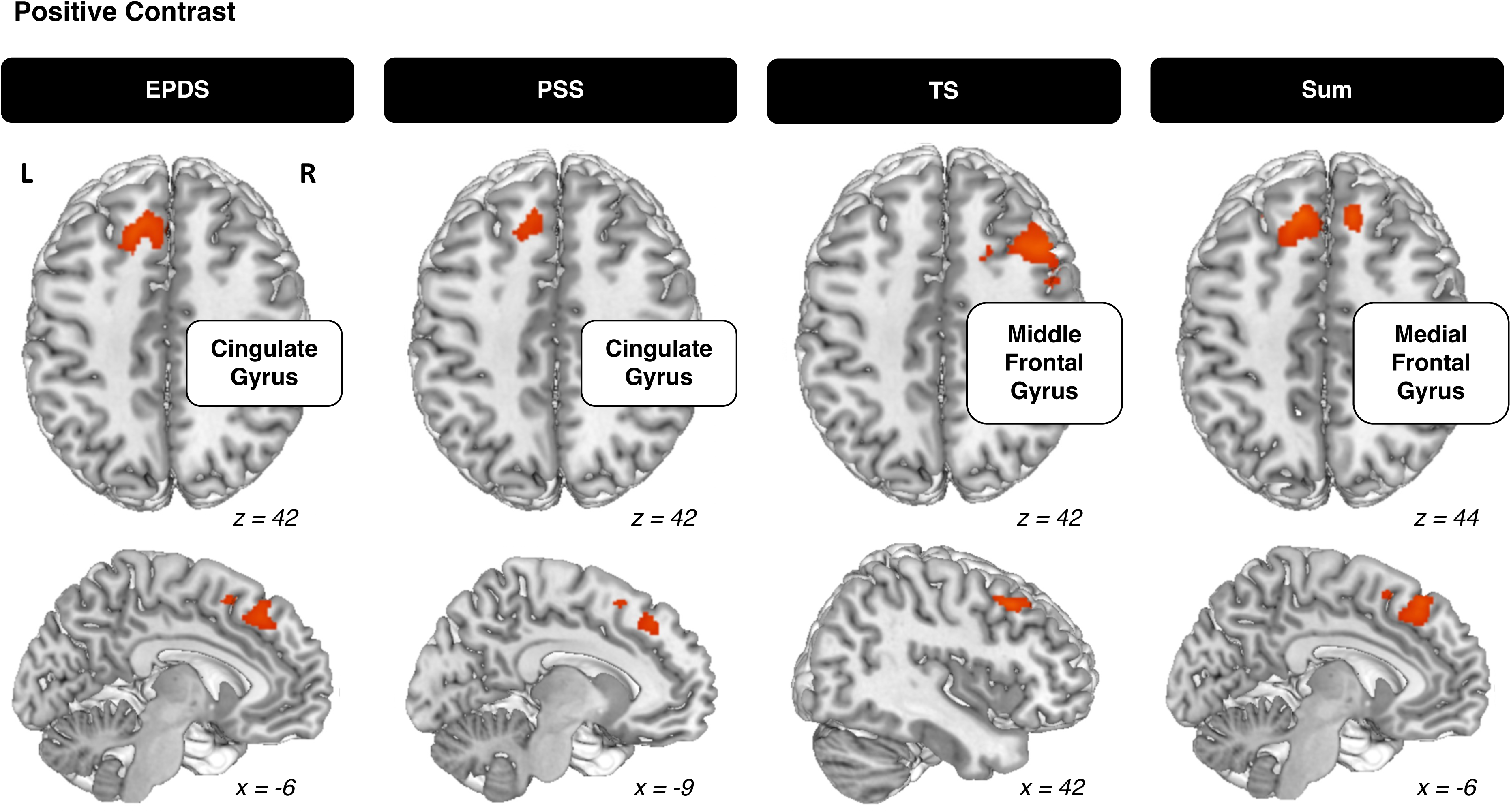
Locations of significant whole-brain activation clusters for No-Go > Go contrast (FDR cluster-level correction of p < 0.05) with maternal psychosocial adversity measures as covariates in the positive contrast. Activation was found through whole-brain regression group analyses (n = 68). Surface renderings of activation patterns were created in Mango (http://ric.uthscsa.edu/mango/) with significant clusters overlayed on Colin27 T1 segmented MNI template. Views shown: superior axial (top) and sagittal right (bottom). L and R represent the left and right sides of the axial surface, respectively. The x and z values represent the position of the respective axes from which the image was taken (x = left/right, z = top/bottom).

## DISCUSSION

In this study, we investigated the functional neuroanatomy of inhibitory control in children 5 to 7 years of age in a low-resource environment. We then examined their relationship with childhood psychosocial adversity (PA) and maternal PA variables, to determine how different forms of PA could explain the activation patterns we observed. As impairments in executive functioning (EF) have been proposed to serve as a mechanism for the relationship between PA and psychopathology, it was necessary to assess how PA is related to inhibitory control, a form of EF, in this environment. We observed significant activation in the inhibitory control contrast (i.e., NoGo versus Go, NGvG) in the bilateral insula, left declive (vermis), right superior frontal gyrus, bilateral superior temporal gyri, and bilateral caudate. In terms of childhood PA, greater neglect was associated with weaker activation in the right posterior cingulate in the NGvG contrast, while greater family conflict and greater household economic stress were related to increased activation in the left medial and right superior frontal gyri, respectively. For exposure to maternal PA, greater maternal stress and depression were all related to increased activation in the NGvG contrast as measured through the EPDS (left cingulate gyrus), PSS (left cingulate gyrus), and TS (right middle frontal gyrus), as well as a summary score of these measures (left medial frontal gyrus). Overall, different forms of PA were associated with unique activation patterns in the NGvG contrast.

The first aim of this study was to determine the regions underlying inhibitory control in the Go/No-Go (GNG) task in a population of children underrepresented in current literature–children 5 to 7 years of age in a low-resource environment. As much of existing literature on developmental cognitive neuroscience relies on populations from Western, educated, industrialized, rich, and democratic areas, there is lacking socio-demographic diversity that is necessary to identify the complexity involved with environmental factors and their relation to neurodevelopment (Garcini et al., 2022). Assessing whether previous results are generalizable to this new population, allows us to understand the extent to which previous explanations in this subject are applicable.

To our knowledge, this study is novel in its usage of fMRI with children in this age group and resource environment. However, the GNG task has been widely used to measure the neural correlates of inhibitory control. Our findings for the inhibitory control contrast (NGvG) were consistent with previous studies examining older children in high resource environments, with strong activation observed in regions related to inhibitory control, particularly in the prefrontal cortex, including the bilateral superior frontal gyri and caudate extending into the bilateral cingulate gyri and right posterior cingulate (McKenna et al., 2017; Houdé et al., 2010).

Large clusters of activation were also observed in the bilateral insula. While this area is not consistently found to be activated in GNG tasks, it has been cited as active during GNG tasks with older children and adolescents due to greater recruitment of attention-based areas in tasks (Houdé et al., 2010; McKenna et al., 2017). When considering the focal patterns of activation as part of the fronto-parietal network, the insula plays an important role in the ventral attention network (Zhang et al., 2017). Recent research has reported the insula to be a “gatekeeper” to various networks important for EF; more broadly, it has been noted that inhibitory control is a cognitive process that involves the connectivity of various networks rather than those solely in the prefrontal area (Molnar-Szakacs & Uddin, 2022; Zhang et al., 2017). The involvement of the ventral attention network is important to consider for the GNG task specifically, as the No-Go stimulus appears less frequently, and requires both a stimulus-driven behavioral response and an individual’s intention-driven response (Cope et al., 2020). Though our age group is not well represented in the literature, it is reasonable that the increased insula activation observed in older children and adolescents would be even greater in younger children who have less well-defined networks and may require even greater attention in such tasks. Additionally, widespread activation outside of prefrontal areas was a finding consistent with our hypothesis, as the prefrontal cortex continues to develop beyond this age, thus refined networks solely within the prefrontal cortex and nearby areas would not be expected in children of this age (Kolk & Rakic, 2022).

The second aim of this study was to understand the relationship between inhibitory control and childhood PA. For our sample, the average sum score of childhood PA based on the CPAS was 39.46 ± 23.32 out of a maximum of 216. This average value is representative of the larger sample as Sullivan et al., who utilized the CPAS for their sample of 154 children from the BEAN project, had an average sum score of 39.33 ± 21.66 (2022). The individual subscale measures also resemble those from Pirazzoli et al., 2022. As this scale was recently developed specifically for this study, we are only able to compare our value with those within the BEAN project. It is important to note that while the individual subscales and total score may not seem very high, even rating “1: Rarely” for an item can indicate exposure to a severe form of adversity in the community (e.g., intimate partner violence, crimes in the community).

On the behavioral level, there were no significant relationships found between task accuracy and presence of PA risks. While studies report mixed results for the behavioral relationship, there is generally an inverse relationship found between inhibitory control and childhood PA (Schäfer et al., 2023). It is possible that no significant relationships were found due to the high accuracy rates with limited variability in our sample. Additionally, because of this similar behavioral performance combined with differing activation patterns, these changes in activation may relate to compensatory strategies, as is discussed further below (Mark et al., 2019; Soloveva et al., 2020).

At the neuroimaging level, significant results were observed when considering the relationship between the childhood PA subscales neglect, family conflict, and household economic stress. We hypothesized that greater childhood PA would be associated with decreased activation in the inhibitory control contrast as previous studies have found decreased activation in the cingulate cortex, middle frontal gyrus, and superior frontal cortex in children with exposure to PA (Bruce et al., 2013; Cará et al., 2019). This hypothesis was supported in the neglect condition, where greater childhood neglect was associated with weaker NGvG activity in the right posterior cingulate. We also hypothesized that in individuals with similar performance, there may be compensatory patterns that relate to increased activation, likely in more posterior areas (Ma et al., 2012; Bruce et al., 2013). For the other significant subscales, greater family conflict and greater household economic stress were associated with increased activation in the medial frontal gyrus (extending to the anterior cingulate) and the superior frontal gyrus (extending to the medial frontal gyrus), respectively. While the posterior cingulate is involved in attention, the medial frontal gyrus, anterior cingulate, and superior frontal gyrus are more directly involved with inhibitory control and decision making (Leech & Sharp, 2014; Talati & Hirsch, 2005; Friedman & Robbins, 2022; du Boisgueheneuc et al., 2006).

There are two possible explanations for these differences in activation patterns according to their related subscales. First, when considering the areas of activation, children with greater childhood PA had increased activation in areas more directly related to inhibitory control (e.g., medial frontal gyrus) and decreased activation in areas less directly related to inhibitory control (posterior cingulate). As previously established, inhibitory control is dependent on the interplay between different networks. Considering there was no significant difference in behavioral performance between these individuals, observing increased activation reflects a form of inefficient recruitment of regions, and may be indicative of modified baselines of activation for substrates of inhibitory control (Farah et al., 2020; Lesnik, 2007; Mark et al., 2019). Some researchers note recruitment of alternate networks/regions to maintain performance while others have found that greater compensatory activation does not necessarily only involve posterior/alternate areas, and can also include task-relevant areas as a result of the greater mental effort required during difficult tasks (Barulli & Stern, 2013; Naccarato et al., 2006; Soloveva et al., 2020). When considering how areas active as part of the fronto-parietal network relating more to decision-making were more active, these activation patterns may be considered a form of compensation.

Another explanation utilizes the deprivation and threat model to understand the impacts of distinct forms of early adversity. While children in areas of high PA frequently have various forms of adversity co-occur, if these aspects are examined separately–between PA factors that relate to deprivation (absence of factors expected for healthy child development) and threat (presence of factors that negatively impact child development)–then unique effects of neurodevelopment can be identified (McLaughlin et al., 2014). When considering childhood adversity related to inhibitory control, deprivation has been reported to have a greater in magnitude reduction in inhibitory control than threat (Johnson et al., 2021). Neglect has commonly been studied as a measure of deprivation, associated with impairments in inhibitory control behaviorally, and with decreased activation in brain networks as a result of the absence of caregiving inputs that traditionally aid in social and cognitive development (Johnson et al., 2021; McLaughlin, 2016). While multiple subscales collected for childhood PA could be categorized as deprivation (neglect, caregiver emotional availability, social isolation), the neglect subscale was the only one identified to have this relationship. For those related to threat (harsh discipline and abuse, physical intimate partner violence, maternal depression, family conflict, household economic stress, community adversity), two subscales were related to increased activation. Threat typically has a larger role related to emotional/stress processing with the hypothalamic-pituitary-adrenal axis and cortisol release, but can also be associated with reduced EF and brain activation due to its relationship with prefrontal cortical circuitry, albeit to a lesser extent than with deprivation (McLaughlin, 2016; Ivy et al., 2010; Merz et al., 2023). The increased activation in task-relevant areas observed in our sample for the threat-related subscales is not reflective of improved EF in these children and can be interpreted as a different presentation of affected patterns of neural networks (Johnson et al., 2021).

As previous research has explored the relationship between the number of adverse childhood experiences (ACEs) related to impacts on EF (e.g., cumulative impact model of ACEs), it was hypothesized that children with greater PA, as measured through the summary score of childhood PA values, would be strongly associated with behavioral and activation differences (McLaughlin et al., 2014). However, in our sample, this was not observed. This absence of finding emphasizes the importance of considering how different types of adversity manifest, rather than relying on a combinatory value, especially as existing literature has found adverse factors related to low-resource environments (neighborhood poverty, lower family income) likely to directly impact brain activation patterns related to inhibitory control, which is associated with psychopathology later in life (Taylor & Barch, 2022; Tomasi & Volkow, 2023; Tomlinson et al., 2020).

The final aim of the study was to determine how maternal PA relates to both the neural and behavioral manifestations of inhibitory control. Previous literature has observed maternal depressive symptoms experienced during early childhood to predict the child’s EF difficulties at later ages, including deficits in inhibitory control (Baker & Kuhn, 2018; Hughes et al., 2013; Wang & Dix, 2017). For our sample, the average maternal stress and depression scores were: 3.00 ± 3.73; PSS: 11.94 ± 8.00; TS: 34.72 ± 9.40; sum: 49.66 ± 17.22. Comparing these values and individual subscale values to larger samples from this study yield similar results (e.g., sum = 52.26 ± 17.23), highlighting the representativeness of our subset to the larger sample (Jensen et al., 2019; Sullivan et al., 2022). When considering how our values compare to those in other areas, the EPDS value resembles those categorized as “non-cases” for depression in a separate Bangladesh sample, while our PSS and TS values are similar in magnitude to those in other areas of South Asia (Gausia et al., 2007; Karasz et al., 2013; Mozumder, 2022). In our sample, similar to the childhood PA measures, there was no significant relationship exhibited between GNG task accuracy and maternal PA variables.

From a neuroimaging perspective, no significant results were observed when considering the inverse relationship between inhibitory control activation and maternal PA. When considering the direct (positive) relationship between inhibitory control activation and maternal PA, greater maternal PA assessed through the EPDS, PSS, TS, and the summary score, were all associated with increased activation in the cingulate gyrus, middle frontal gyrus, and medial frontal gyrus. It is challenging to integrate these findings with previous literature, as there have been mixed results related to the relationship between maternal PA and inhibitory control. For example, while Farah et al. (2020) observed that children with higher maternal PA have decreased functional connectivity between areas related to imagination and auditory processing, Demir-Lira et al. (2016) found greater regional homogeneity in resting-state fMRI in the left prefrontal cortex in young children with higher early life stress. Meanwhile, Pozzi et al. (2021) reported no association between negative maternal behavior and differences in functional connectivity patterns in children. As all maternal PA variables were associated with increased activation in areas directly related to the inhibitory control task, this is possible to be explained with what some researchers predict in relation to compensation, as there was diffuse increased activation resulting in the maintenance of behavioral performance (Naccarato et al., 2006; Soloveva et al., 2020). Contextualizing these measures within the deprivation and threat model is also difficult as maternal stress and depression involve both threat and deprivation effects, as they not only involve the addition of stress–a negative environmental factor (threat)–but also lead to less warm and responsive parenting behaviors for the child (deprivation) (Henry et al., 2020). Regardless, the presence of increased activation reflects neural correlates of inhibitory control that differ from previous literature, again highlighting the use of varied recruitment of brain regions for inhibitory control in this novel population.

### Limitations

Though this study successfully employed neuroimaging techniques to examine the relationship between PA and children’s inhibitory control in a sample of children growing up in profound poverty, there were some limitations. First, as these techniques relied on assessing this relationship through correlational analyses, no information regarding the causality of this mechanism can be stated. Regarding the PA behavioral assessments, though the assessments were adapted to a Bangladeshi context, it is possible that these measures were not fully comprehensive to assess PA and outcomes in this sample. These measures also relied on the assessment of negative psychosocial factors and did not assess the prevalence of protective factors that may have allowed for resiliency within this population. Additionally, while the GNG task utilized in this study has been widely used in conjunction with neuroimaging tools to inform of the neural correlates of inhibitory control, it is possible that this task may not have been sensitive enough, or our sample size may not have been large enough, to detect individual behavioral variance related to performance in this environment. However, in a previous study that utilized a larger sample from the BEAN project, Sullivan et al. (2022) found behavioral differences between participants through a similar GNG task, which indicates that the behavioral results in the present study are likely reflective of the true variance, especially as our sample’s measures were found to be representative of this larger sample. It is also important to note that despite the wide use of the GNG task to interpret the contrast between behavioral variables and inhibitory control through successful inhibitions (NoGo > Go), this approach has not been without criticism, with findings suggesting that error-related contrasts provide better information (Paige et al., 2024; Weigard et al., 2020). While these studies focused on older and White populations, future studies should work to determine which contrast from the GNG task is most closely associated with social and behavioral measures in more diverse populations. Finally, as there was no middle or high resource group from Bangladesh to compare the children in our sample to, it is difficult to know whether the differences observed in the activation patterns, as well as the diffuse clusters, would also be revealed in other groups living in this area. Despite these limitations, examining brain function underlying GNG in children growing up in a low-resource environment contributes to a more comprehensive image of the neural activation patterns present in the development of inhibitory control.

## Conclusion

In conclusion, this study presents results from neuroimaging data that explores the relationship and identifies the neural correlates of inhibitory control and PA in children within a low-resource environment, providing greater cultural and contextual diversity to this subject area. Results indicate decreased activation related to childhood neglect and increased activation related to greater childhood, household, and maternal stress and depression. These results are reflective of modified neural substrates for inhibitory control and align with proposed explanations of compensatory mechanisms and differences in effects between threat and deprivation adverse factors. The results emphasize the importance of conducting research in low-resource environments to successfully identify how inhibitory control may serve as a mechanism for the relationship between early life adversity and psychopathology. Future studies should work to assess older children as well as adults in similar low-resource environments, through both cross-sectional and longitudinal methods, to establish typical and atypical inhibitory control activation patterns in these areas. As this present study has shown measures of PA relate to differences in inhibitory control activation patterns, future studies could utilize interventions that work to minimize PA and see if there are observable impacts in the regions underlying inhibitory control, and for which early years the brain is most amenable to restoring impaired EF.

