## Supplementary Material for "Examining the relationship between psychosocial adversity and inhibitory control: an fMRI study of children growing up in extreme poverty"

***Table 1**: MRI scan parameters*

| **Structural T1-weighted MPRAGE[1] images:** | **Gradient-echo EPI[2] images:** |
|---|---|
| TR = 2500 ms<br>TE = 3.47 ms<br>176 sagittal slices<br>1 $mm^3$ voxels<br>FOV = 256 mm | TR = 2000 ms<br>TE = 3.47 ms<br>32 axial slices (interleaved)<br>Voxel dimensions 3 x 3 x 4.8 mm<br>In-plane resolution 64 x 64<br><br>132 functional BOLD volumes |

[1]MPRAGE = Magnetization Prepared - RApid Gradient Echo
[2]EPI = echo-planar imaging

### ***Table 2:*** *Correlational analyses*

| | *Age* | *Go Accuracy* | *NoGo Accuracy* | *CPAS: Neglect* | *CPAS: Emotional Availability* | *CPAS: Harsh Discipline + Abuse* | *CPAS: Social Isolation* | *CPAS: Physical Intimate Partner Violence* | *CPAS: Family Conflict* | *CPAS: Depression* | *CPAS: Household Economic Stress* | *CPAS: Community Adversity* | *CPAS: Sum* | *EPDS* | *PSS* | *TS* | *Maternal Health Sum* |
|---|---|---|---|---|---|---|---|---|---|---|---|---|---|---|---|---|---|
| *Age* | 1 | | | | | | | | | | | | | | | | |
| *Go Accuracy* | 0.2 | 1 | | | | | | | | | | | | | | | |
| *NoGo Accuracy* | -0.1 | 0.16 | 1 | | | | | | | | | | | | | | |
| *CPAS – Neglect* | 0.19 | 0.18 | 0.06 | 1 | | | | | | | | | | | | | |
| *CPAS: Emotional Availability* | 0.14 | 0.01 | 0.02 | 0.24 | 1 | | | | | | | | | | | | |
| *CPAS: Harsh Discipline + Abuse* | 0.11 | 0.10 | -0.18 | 0.32 | -0.06 | 1 | | | | | | | | | | | |
| *CPAS: Social Isolation* | 0.25 | 0.08 | 0.08 | 0.45 | 0.08 | 0.41 | 1 | | | | | | | | | | |
| *CPAS: Physical Intimate Partner Violence* | 0.06 | 0.09 | 0.05 | 0.24 | 0.26 | 0.20 | 0.53 | 1 | | | | | | | | | |
| *CPAS: Family Conflict* | 0.22 | -0.01 | 0.04 | 0.34 | 0.17 | 0.30 | 0.78 | 0.56 | 1 | | | | | | | | |
| *CPAS: Depression* | 0.19 | 0.07 | 0.09 | 0.41 | -0.01 | 0.35 | 0.74 | 0.38 | 0.63 | 1 | | | | | | | |
| *CPAS: Household Economic Stress* | 0.16 | 0.22 | 0.10 | 0.63 | 0.03 | 0.28 | 0.45 | 0.28 | 0.41 | 0.57 | 1 | | | | | | |
| *CPAS: Community Adversity* | 0.16 | 0.12 | 0.00 | 0.34 | 0.15 | 0.28 | 0.31 | 0.42 | 0.33 | 0.34 | 0.42 | 1 | | | | | |
| *CPAS: Sum* | 0.25 | 0.14 | 0.03 | 0.65 | 0.23 | 0.59 | 0.84 | 0.61 | 0.77 | 0.81 | 0.70 | 0.59 | 1 | | | | |
| *EPDS* | 0.14 | 0.10 | 0.01 | 0.25 | 0.12 | 0.23 | 0.46 | 0.23 | 0.30 | 0.41 | 0.36 | 0.12 | 0.44 | 1 | | | |
| *PSS* | -0.01 | -0.01 | 0.05 | 0.14 | 0.27 | 0.20 | 0.48 | 0.16 | 0.34 | 0.38 | 0.34 | 0.13 | 0.43 | 0.78 | 1 | | |
| *TS* | 0.11 | 0.28 | 0.08 | 0.37 | 0.14 | 0.23 | 0.18 | 0.18 | 0.17 | 0.31 | 0.58 | 0.30 | 0.42 | 0.47 | 0.34 | 1 | |
| *Maternal Health Sum* | 0.09 | 0.17 | 0.07 | 0.33 | 0.23 | 0.27 | 0.42 | 0.23 | 0.32 | 0.44 | 0.55 | 0.25 | 0.52 | 0.83 | 0.82 | 0.81 | 1 |
